# Comprehensive Assessment of Ischemic Stroke in Nonhuman Primates: Neuroimaging, Behavioral, and Serum Proteomic Analysis

**DOI:** 10.1101/2023.12.20.572532

**Authors:** Ge Li, Lan Lan, Tingting He, Zheng Tang, Shuhua Liu, Yunfeng Li, Zhongqiang Huang, Yalun Guan, Xuejiao Li, Yu Zhang, Hsin-Yi Lai

**Author notes:** Co-first author (Ge Li, Lan Lan). Correspondence: Yu Zhang, Hsin-Yi Lai Hsin-Yi Lai Yu Zhang.

## Abstract

Ischemic strokes, prevalence and impactful, underscore the necessity of advanced research models closely resembling human physiology. O integrating n ur study in nonhuman primates (NHPs) offers a comprehensive exploration of ischemic stroke, integrating neuroimaging data, behavioral outcomes, and serum proteomics to elucidate the complex interplay of factors involved in stroke pathophysiology. We observed a consistent pattern in infarct volume, peaking at 1-month post-middle cerebral artery occlusion (MCAO) and stabilizing thereafter. This trend was closely correlated with notable changes in motor function and working memory performance. Using diffusion tensor imaging (DTI), we detected significant alterations in fractional anisotropy (FA) and mean diffusivity (MD) values, indicative of microstructural changes in the brain. These findings were strongly correlated with the observed neurological and cognitive deficits, highlighting the sensitivity of DTI metrics in stroke assessment. Behaviorally, the monkeys exhibited a reliance on their unaffected limb for compensatory movements, a response commonly observed in stroke impairment. This adaptation, alongside the consistent findings in DTI metrics, suggests a substantial impact of stroke on motor function and spatial perception. Proteomic analysis through MS/MS functional enrichment revealed two distinct groups of proteins with significant changes post-MCAO. Notably, MMP9, THBS1, MB, PFN1, and YWHAZ emerged as potential biomarkers and therapeutic targets in ischemic stroke. Our findings underscore the complex nature of stroke and the potential of an integrated approach, combining neuroimaging, behavioral studies, and proteomics, for advancing our understanding and treatment of this condition.

## 1. Introduction

Stroke poses a major global health challenge, being the primary cause of long-term disability and the second leading cause of death internationally. Ischemic strokes, which make up about 77% of cases, arise from reduced cerebral blood flow due to arterial blockages, leading to tissue infarction and extensive cellular damage (GBD 2016 Lifetime Risk of Stroke Collaborators, 2018; Narayan et al., 2021). The consequences of stroke are significant, manifesting as neurological impairments such as contralateral hemiparesis, which profoundly affect daily life. The severity and type of these impairments vary with the affected brain areas, often resulting in motor deficits. (Langhorne et al., 2009; Bai et al., 2014). This impacts over 80% of stroke survivors, particularly in upper limb movement, drastically impacting their quality of life (Jones, 2017).

Animal models, especially non-human primates (NHPs), are crucial in stroke research for developing effective treatments. While rodent models have been instrumental, their physiological and anatomical disparities from humans limit the translating potential of research findings (Taha et al., 2022). NHP models, with their closer genetic and physiological resemblance to humans, offer a more accurate representation of human stroke pathology and recovery processes (Fisher et al., 2009; Sommer, 2017; Marshall et al., 2001). These models are particularly advantageous for long-term neuroimaging and behavioral assessments, providing a comprehensive view of stroke’s impact over time (Yi et al., 2017).

Advancements in neuroimaging, particularly diffusion tensor imaging (DTI), have greatly enhanced our understanding of stroke. DTI allows in vivo assessment of tissue integrity and connectivity, providing potential biomarkers for disease progression (Shen et al., 2022; Shaheen et al., 2022). It has been effective in tracking changes in white and grey matter during stroke in both rodent (Jung et al., 2017) and human (Li et al., 2022) studies. DTI metrics, such as fractional anisotropy (FA) and mean diffusivity (MD), provide insights into the microstructural integrity of neural tissue (Hagmann et al., 2006Kronlage et al., 2018), with lower FA values linked to poorer motor outcomes in stroke patients (Okamoto et al., 2021). Chronic white matter injury manifests as increased MD and decreased FA have been consistently observed in strokes patients (Pasi et al., 2016) and NHP stroke model (Won et al., 2020).

Proteomics has emerged as a vital tool in stroke research, identifying biomarkers and therapeutic targets and shedding light on proteomic alterations during various stroke phases (Chen et al., 2022). Global proteomic changes in ischemic core and penumbra regions have been extensively studied in rodent and NHP models, revealing over 7000 differentially expressed proteins implicated in critical processes like cell death, survival, and recovery (Hochrainer & Yang, 2022). Serum proteomics in NHP stroke models provides insights into cellular and biochemical pathways involved in stroke progression from the acute to chronic phases in rodents (Sironi et al., 2001) and NHPs (Stevens et al., 2019).

This study aimed to explore the intricate stages of stroke in NHP stroke models by examining brain microstructural changes, pathological indices, neurological alterations, proteomic changes. We hypothesize that the disruption of blood supply, coupled with neuronal cell damage and white matter integrity impairment, leads to cerebral edema, dysfunction, and structural abnormalities. Furthermore, we anticipate that motor injury symptoms are associated with alterations in DTI metrics, warranting an in-depth exploration of the correlations between DTI parameters and neurological features. Through serum proteomic mapping, we aim to investigate the parallels between NHP stroke models and human stroke, focusing particularly on biomarkers reflective of brain microstructural integrity and their association with the pathophysiological mechanisms of stroke.

## Results

We meticulously conducted middle cerebral artery occlusion (MCAO) to induce ischemic stroke models in non-human primates. This approach involved the integration of neuroimaging data, behavioral assessments, and detailed serum proteomic analysis at both 1-month and 3-month post-MCAO. Our objective was to unravel the intricate interplay of various factors influencing stroke pathophysiology, providing a more holistic understanding of the condition.

### Changes in Infarct Volume and Microstructure in the Brain

Our longitudinal monitoring of the infarct volume using T2-weighted images revealed that the ischemic lesion reached its maximal expansion at 1-month post-MCAO (*p*=0.0146), as shown in **Figure 1A**. Total infarct volume remained consistent between the 1-month and 3-month intervals (*p*=0.9554), suggesting no significant reduction or recovery in lesion size during this period. The infarct area, marked in red on each axial MRI image, varied in location across subjects. This variation is depicted in Figure 1A, which aligns MRI findings with Paxinos Macaque Atlas (Calabrese et al., 2015). Observed lesions encompassed several critical regions, including the orbitofrontal cortex, insular cortex, and components of the basal ganglia such as the putamen, caudate nucleus, olfactory tubercle, and globus pallidus, extending to adjacent white matter regions. The intra-class correlation coefficient for infarct volume evaluations between two independent readers was exceptionally high at 0.998, indicating robust agreement and reliability in these measurements.

**Figure 1.**
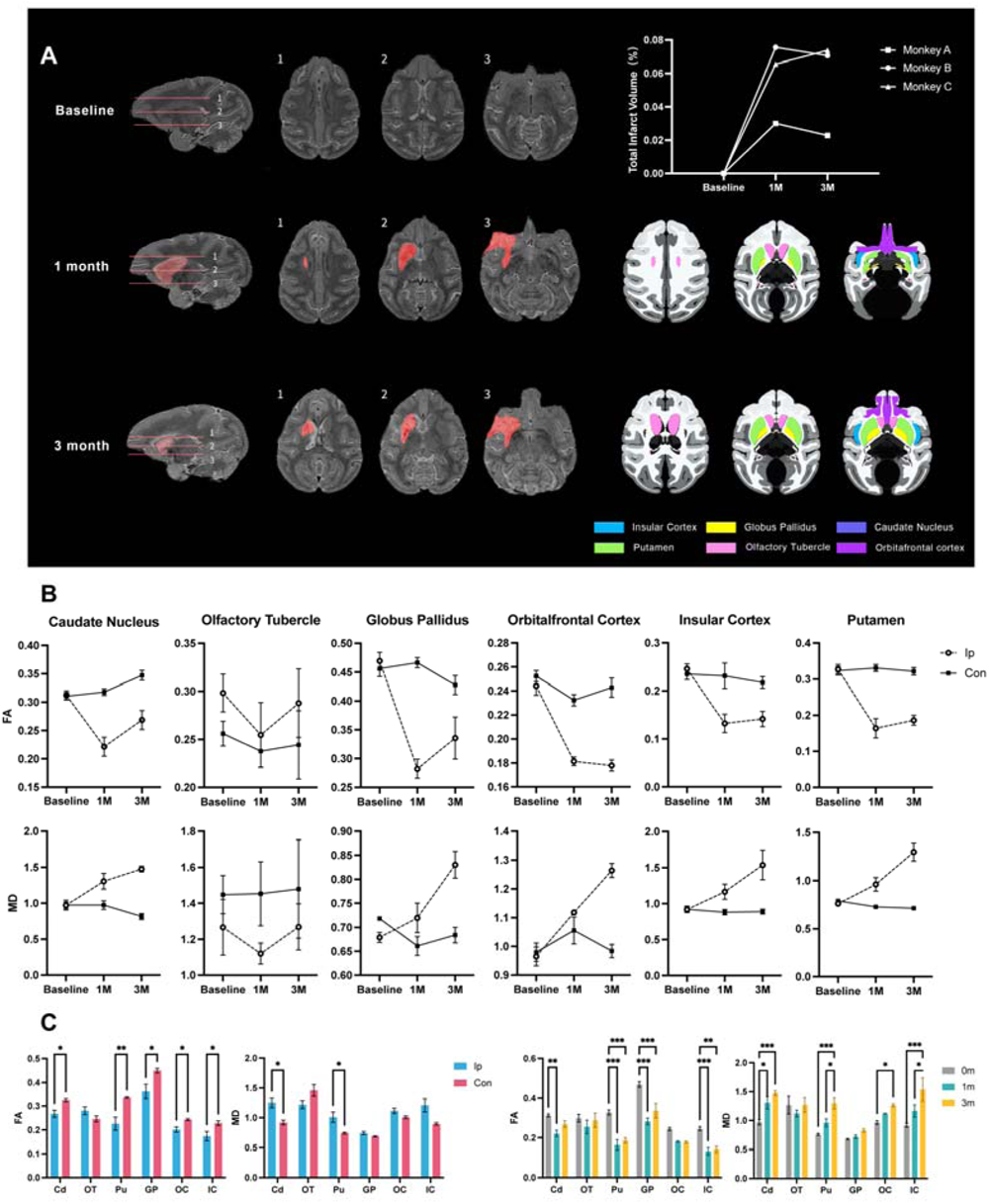
MRI image analysis. (A) T2-weighted images display the ischemic lesion (marked in red) against macaque brain atlas sections. The infarct area expands notably at 1-month post-MCAO, followed by a noticeable reduction by 3-month. (B) In the lesioned left hemisphere, ipsilateral fractional anisotropy (FA) values decrease at 1-month post-MCAO and partially recover by 3-month, but remain reduced compared to the contralateral side. Mean diffusivity (MD) values show an increasing trend over the 3-month period. (C) This segment contrasts FA and MD values between ipsilateral and contralateral regions across six ROIs: caudate nucleus (Cd), olfactory tubercle (OT), putamen (Pu), globus pallidus (GP), orbitofrontal cortex (OC), and insular cortex (IC), measured at baseline, 1-month, and 3-month post-MCAO. Statistical significance levels are indicated as: *** (P<0.001), ** (P<0.01), * (P<0.05). Data are presented as Mean ± SEM, with ’Con’ for contralateral and ’IP’ for ipsilateral.

Both FA and MD values, important metric derived from DTI, were used to evaluate the microstructure alterations resulting from ischemic lesion in the brain, as shown in **Figure 1B**. FA values provide information about the integrity and directionality of white matter tracts in the brain, and MD values measure the average rate of water diffusion within a tissue, regardless of its direction. In general, reduced FA and increased MD values are often observed in areas of brain pathology.

In the ipsilateral regions of interest (ROIs), lesion sites, FA values showed a significant decline at 1-month post-MCAO compared to baseline, indicating disrupted white matter integrity. Interestingly, these values began to recover beyond the 3-month post-MCAO. Across all ipsilateral ROIS, FA values were significantly lower than those on the contralateral side (p < 0.05), excepting in the olfactory tubercle. Notable changes in FA were observed in the caudate nucleus, putamen, globus pallidus, and insular cortex, with significant differences between baseline and the 3-month post-MCAO, as shown in **Figure 1C**. In contrast, MD values displayed a consistent increase in ipsilateral ROIs, lesion sites, from baseline to 3-month post-MCAO, suggesting increased water content and breakdown of normal cellular structures. At 3-month post-MCAO, significant increases in MD were shown in the caudate nucleus, putamen, orbitofrontal cortex, and insular cortex compared to baseline (Figure 1C). The AD and RD are other two specific metrics derived from DTI. Alterations in AD values are indicative of changes in axonal integrity, while variations in RD values correspond to the status of myelination sheaths within the brain’s white matter. Detailed result on these metrics and their implications were provided in the Supplemental Materials (**Figure S1**).

### Behavioral Outcomes

Comprehensive behavioral assessment in nonhuman primates were conducted before MCAO induction, and at 1-month and 3-month post-MCAO, including neurologic deficit evaluation, motor function analysis (hill and valley staircase task, HVST), working memory assessment (delayed-response task, DRT), and cognitive function evaluation (object-retrieval detour task, ORDT). Before MCAO, monkeys exhibited normal neurologic functions with average score of 6.00±1.63, as shown in Figure 2A. Significant neurological deficits, including right limb paralysis and reduced tone, were observed at 1-month post-MCAO, with scores rising to 29.67±0.94 (*p*<0.0001). No significant improvement was noted at 3-month post-MCAO (*p*=0.2302, average score of 28.33±0.94 for 3-month post-MCAO). Figure 2B shows the results of hill and valley staircase task (HVST). Before MCAO, all monkeys almost completed hill and valley staircase tasks using both upper limbs. At 1-month and 3-month post-MCAO, impaired right upper limb function hindered task performance, whereas left limb function remained unaffected. In the valley staircases task, all monkeys effectively used their left upper limb to retrieve food from the left staircase. Working memory was evaluated by the delayed-response task (DRT), as shown in Figure 2C. Initially, monkeys had an average correct response rate of 85.00±17.80 in the DRT. This rate significantly dropped to 63.33±22.11 at 1-month post-MCAO (*p*=0.008), indicating a decline in working memory, with no further deterioration at 3-month (61.67±17.95, *p*=0.8283). The cognitive function was assessed by the object retrieval detour task (ORDT). Before MCAO, the success rate was 91.04±6.92%, with barrier hits of 1.73±1.69, as shown in Figure 2D. A significant drop in success rate (81.04±10.97%, *p*=0.0075) and an increase in barrier hits (4.20±4.78, *p*=0.0793) were observed at 1-month post-MCAO, indicating cognitive impairment. By 3-month, both success rate (93.75±4.11, *p*=0.0004) and barrier hits (0.67±1.07, *p*=0.0116) showed significant improvement, aligning closely with pre-surgery performance.

**Figure 2.**
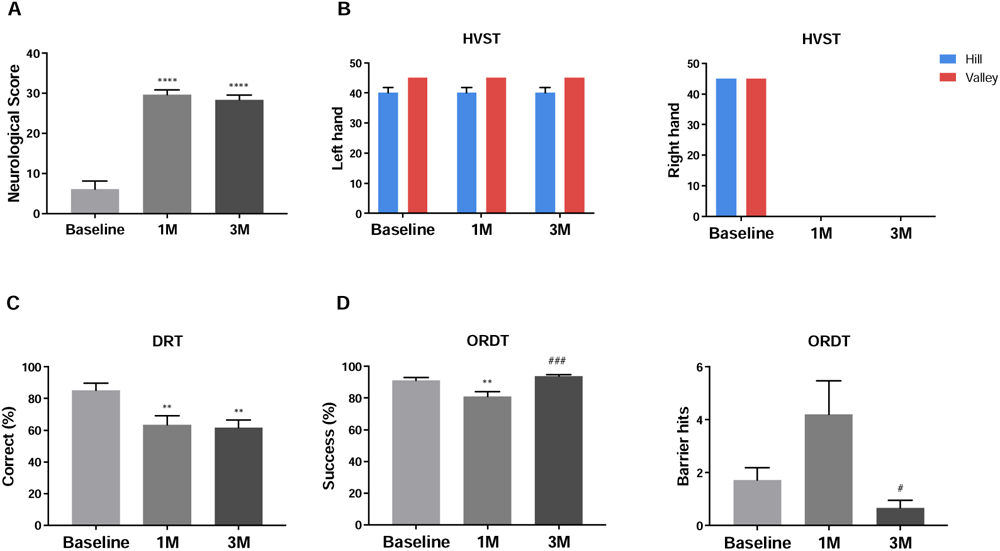
Behavioral Assessment. (A) Neurologic score showed a significant increase at 1-month and 3-month post-MCAO compared to the baseline. (B) Motor function in the left upper limb remained unaffected in both hill and valley staircases tasks (HVST) post-MCAO, maintaining baseline levels. Conversely, right upper limb was notably impaired post-MCAO, reflecting a decline from baseline performance. (C) In the delayed-response task (DRT), a significant decrease in the average correct response rate was observed at both at 1-month and 3-month post-MCAO. (D) The object retrieval detour task (ORDT) showed dynamic changes post-MCAO: the success rate of first-attempt reward retrieval significantly decreased at 1-month but recovered by 3-month, while the frequency of barrier hits increased initially at 1-month post-MCAO, indicating increased attempts to reach the reward through the transparent wall, and then showed marked improvement by 3-month. Data present Mean ± SEM of from 3 monkeys across 5 independent experiments. Significance denoted as: *** (P<0.001), ** (P<0.01), * (P<0.05) versus baseline, and ### (P<0.001) versus 1-month, all determined by Student’s t-test.

### Serum Protein Identification and Quantitation

Using liquid chromatography-mass spectrometry (LC-MS)/MS, we analyzed the tryptic peptides from serum protein samples collected before MCAO, and at 1-month and 3-month post-MCAO. A total of 5214 peptides and 799 proteins were identified. Proteins with a fold change greater than 1.5 and a *p*-value < 0.05 (Student’s t-test) were considered differentially expressed. Post-MCAO, 60 proteins at 1-month and 91 proteins at 3-month showed significant changes in expression, and 30 proteins significantly altered at 3-month post-MCAO compared to 1-month post-MCAO, as shown in Figure 3A. An hierarchical cluster analysis was performed to visualize these changes (see details in **Figure S2**), Using the Mfuzz package, 6 distinct clusters were identified based on differential expression criteria (fold change > 1.5, *p*-value < 0.05), meantime, to gain further insights into the biological implications of these clusters, Gene Ontology (GO) and Kyoto Encyclopedia of Genes and Genomes (KEGG) enrichment analyses were conducted as shown in Figure 3B. Applying a stricter fold change threshold greater than 5.0 and *p-*value less than 0.05, 42 candidate proteins were identified across clusters (see details in Table S1), with notable upregulation in Cluster 1 and Cluster 2. Notable proteins involved in cell morphogenesis and migration included actin beta like 2 (ACTBL2), actin gamma 1 (ACTG1), f-actin-capping protein subunit alpha-2 (CAPZA2), profilin 1 (PFN1), tropomyosin 4 (TPM4), moesin (MSN), WD repeat domain 1 (WDR1), S100 calcium binding protein A4 (S100A4), monoglyceride lipase (MGLL), coactosin like f-actin binding protein 1 (COTL1), Ras suppressor protein 1 (RSU1), parvin beta (PARVB), and Transgelin (TAGLN2). Proteins like matrix metallopeptidase 9 (MMP9), thrombospondin 1 (THBS1) and heat shock protein HSP 90-alpha (HSP90AA1) in Cluster 1, and L-lactate dehydrogenase A chain (LDHA), actinin alpha 1 (ACTN1), coronin 1A (CORO1A), coronin 1B (CORO1B), glutathione S-transferase pi 1 (GSTP1), and tyrosine 3-monooxygenase/tryptophan 5-monooxygenase activation protein zeta (YWHAZ) in Cluster 2 were identified as key in apoptotic regulation, were also detected. Myoglobin (MB) in Cluster 1 and Serpin family B member 1 (SERPINB1), associated with coagulation regulation, were also detected. To further understand the intricate relationships and functional dynamics among the proteins identified in clusters 1 and 2, we employed the STRING database to construct a protein-protein interaction network, as shown in Figure 3C. This approach enabled us to reveal potential interconnections and explore the broader functional implications of these proteins in the context of ischemic stroke pathophysiology.

**Figure 3.**
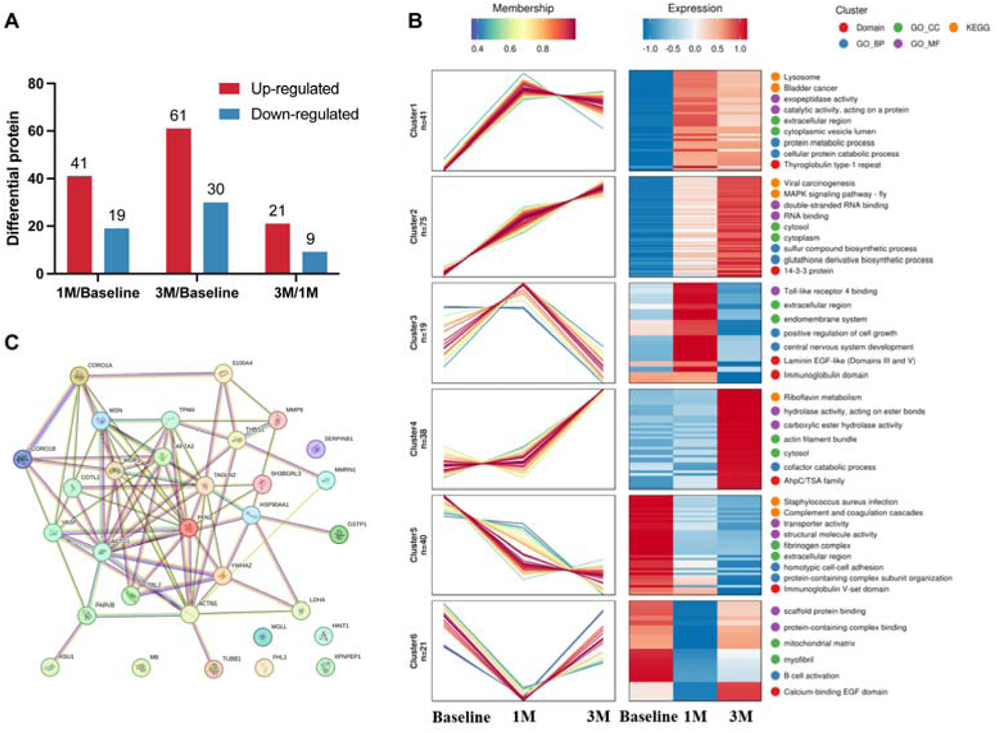
Serum protein expression analysis. (A) Comparative graph depicting the count of differentially expressed proteins before MCAO and at 1-month and 3-month post-MCAO, highlighting temporal protein expression changes. (B) Application of the Mfuzz package for hierarchical cluster analysis identified 6 distinct clusters based on a fold change > 1.5 and *p*-value < 0.05. (C) Visualization of the protein-protein interaction network for proteins in Cluster 1 and Cluster 2, utilizing the STRING database to elucidate potential interconnections and functional relationships.

### Correlations between MRI data, behavioral outcomes, and serum proteomics

Correlation analysis between MRI data, behavioral outcomes, and serum proteomics were performed in Figure 4. The relationships between infarct volume and behavioral outcomes were illustrated in Figure 4A. Volume analysis uncovered a significant positive correlation existed between infarct volume and neurological scores (*p*=0.0004). Conversely, a notable inverse correlation was observed between infarct volume and the accuracy of responses in the DRT correct (*p*=0.0123). However, no significant correlation emerged in relation to the ORDT scores. These findings suggest that the infarct volume is associated with alterations in neurological and working memory functions. Figure 4B showed the correlations between DTI metrics and behavioral outcomes. FA values positively correlated with DRT correct (*p*=0.0092), while negative correlations were observed with MD (*p*=0.0270), AD (*p*=0.0459) and RD (*p*=0.0105) values. Neurological score negative correlated with FA (*p*=0.0024), but no significant correlations were identified with ORDT outcomes. Additionally, Cluster 1 and Cluster 2 negatively correlated with FA (*p*=0.0433) and positively with MD (Cluster 1: *p*=0.0061, Cluster 2: *p*=0.0061), AD (Cluster 1: *p*=0.0045, Cluster 2: *p*=0.0311), and RD (Cluster 1: *p*=0.0045, Cluster 2: *p*=0.0031), as shown in Figure 4C. Figure 4D showed both clusters showed a negative correlation with DRT correct (Cluster 1: *p*=0.0165, Cluster 2: *p*=0.0035) and a positive correlation with neurological function scores (Cluster 1: *p*=0.0118, Cluster 2: *p*=0.0160)), indicating more severe deficits with higher cluster values. In addition, correlations between DTI metrics and Serum Protein were showed in **Table 1**. FA values negatively correlated with proteins in Cluster 1, including MMP9, THBS1, MB, and MD values showed positive correlation with proteins in Cluster 2, including SERPINB1, ACTBL2, ACTG1, RUS1, MGLL, PFN1, WDR1, TAGLN2, YWHAZ, ACTN1.

**Figure 4.**
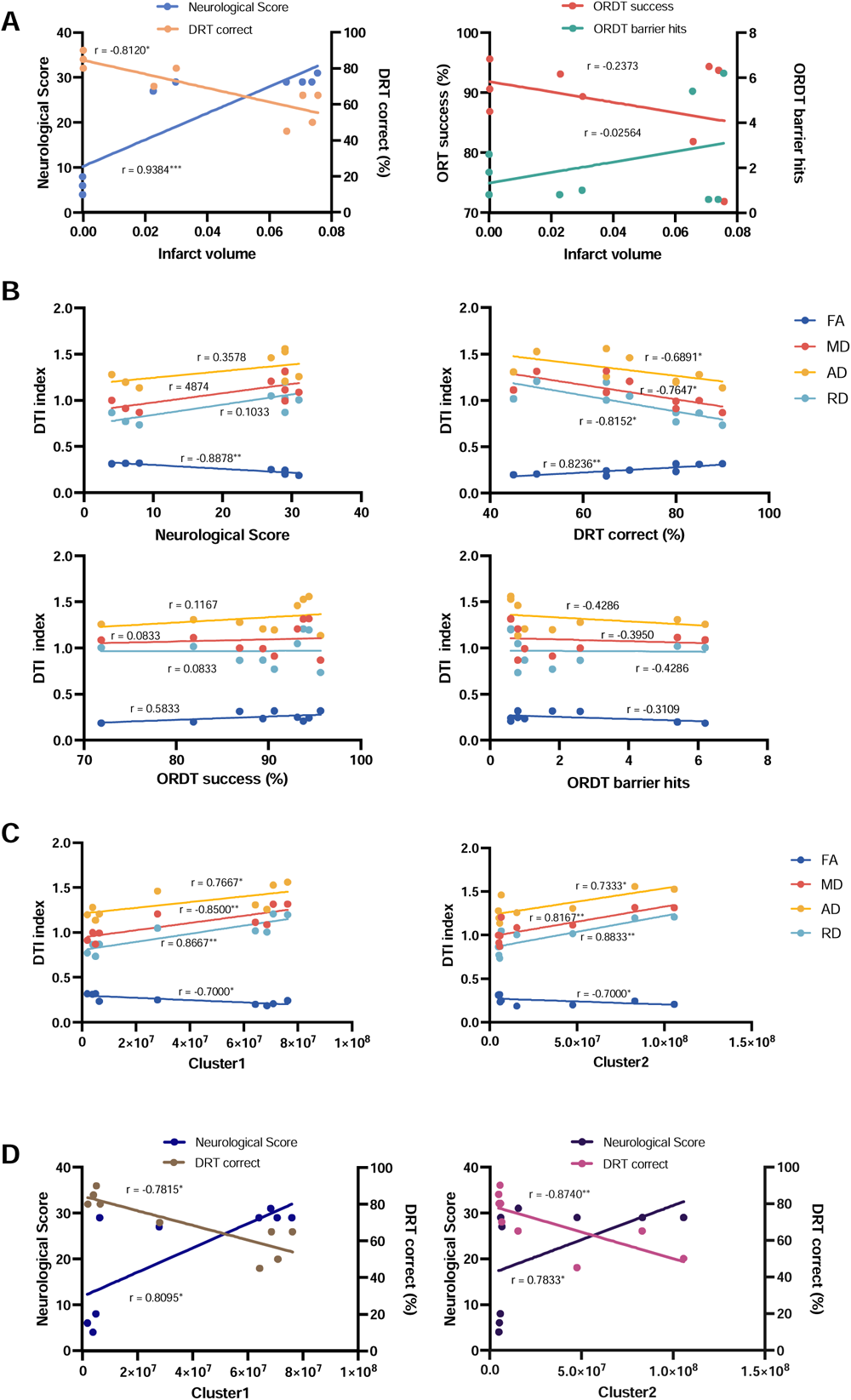
Correlations between infarct volume, DTI metrics, behavioral outcomes, and serum proteomics. (A) A negative correlation between infarct volume and delayed-response Task (DRT) correct. A positive correlation with neurological score, and no significant correlations with object retrieval detour task (ORDT) success (*p*=0.5422) or barrier hits (*p*=0.9545). (B) Detail the correlations between DTI metrics, including fractional anisotropy (FA), mean diffusivity (MD), axial diffusivity (AD) and radical diffusivity (RD), and behavioral outcomes: FA shows a significant positive correlation with both DRT correct and neurological scores (*p*=0.0024), while MD, AD, and RD exhibit negative correlations with DRT correct. In contrast, no significant correlations are observed between any of the DTI metrics and ORDT success (FA, *p*=0.1080, MD, *p*=0.8432, AD, *p*=0.7756, RD, *p*=0.8432) and ORDT barrier hits (FA, *p*=0.4121, MD, *p*=0.2922, AD, *p*=0.2502, RD, *p*=0.2502). (C) Explore the correlation between serum proteomic clusters (Cluster 1 and Cluster 2) and DTI metrics, with both clusters showing significant correlations with each DTI metric. (D) Cluster 1 is negatively correlated with DRT correct and positively correlated with neurological scores. Cluster 2 exhibits a similar pattern, showing a negative correlation with DRT accuracy and a positive correlation with neurological scores. Significance levels are indicated as: *** (*p*<0.001), ** (*p*<0.01), * (*p*<0.05).

**Table 1.**
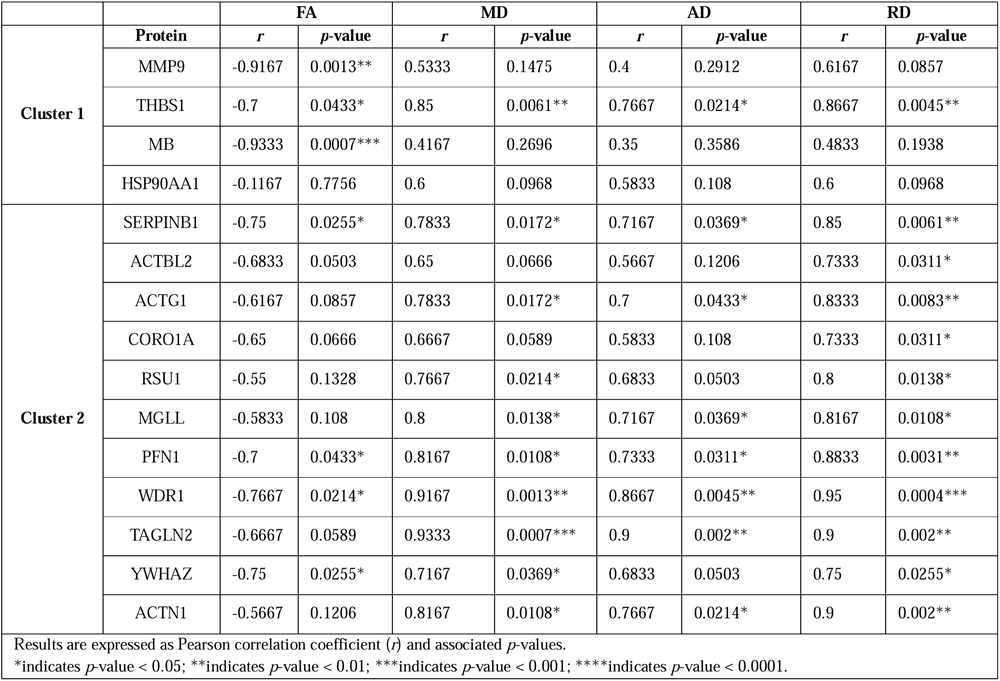
Correlation between DTI metrics and Serum Protein.

## Discussion

In this study, we conducted a detailed analysis of MRI data, behavioral outcomes, and serum proteomics in a non-human primate (NHP) ischemic stroke model. Our study focused on the intricate relationship between neuroimaging findings, behavioral functions, and proteomic profiles following an ischemic stroke, aiming to integrate these diverse data streams for a holistic understanding of stroke pathophysiology. We observed the infarct volume peaked at 1-month post-MCAO and remained constant up to three months, a trend that correlated with motor function changes, neurological impairment, and working memory performance. Additionally, we found that the decline in FA and the increase in MD were significantly correlated with both motor and cognitive deficits. The serum proteomic analysis, integrating protein identification and quantitation, revealed critical correlations with neuroimaging and behavioral findings. These findings open new avenues for a multimodal approach to understanding stroke pathology and recovery, integrating neuroimaging, behavioral assessment, and molecular profiling.

Our findings revealed that the infarct volume reached its maximum size one month after the MCAO procedure and then stabilized, showing no significant change for up to three months (Figure 1A). The structural brain changes during stroke’s acute phase, characterized by infarct lesions and loss of manual dexterity, are possibly due to oxidative stress and other factors induced by reperfusion injury (Zweier & Talukder, 2006). These changes correspond with our observed correlations between injury volume and neurological function assessments (Figure 4A). In addition, we detected a significant decrease in FA and an increase in MD across various brain regions, compared to the contralateral side of the injury area (Figure 1B). This aligns with prior studies linking DTI metrics to injury patterns in motor fiber bundles and motor dysfunction (Ma et al., 2014; Kwak et al., 2010). The changes in FA and MD around the infarcted areas suggest white matter damage, impacting neurological function and recovery. Interestingly, FA showed signs of recovery at 3-month post-MCAO, while MD continued to increase, suggesting early-stage tissue repair and axonal regrowth, alongside ongoing tissue damage or inflammation (Figure 1B). This asynchronization may be related to the specificity of the damage and recovery of white matter microstructure. Our findings corroborate previous studies indicating sensitivity of FA in predicting behavioral outcomes and its potential as a clinical tool for assessing stroke prognosis (Lindenberg et al., 2012) (Okamoto et al., 2021).

In the behavioral aspect, one month following the MCAO procedure, the monkeys in our study demonstrated notable neurological deficits (Figure 2), including paralysis and reduced muscle tone in their right limbs, which persisted without significant improvement over the subsequent three months. This reflects the substantial impact of stroke and correlates with research linking more severe motor deficits and reaction time delays to greater ipsilateral hemispheric damage (Foltys et al., 2003). DTI metrics like FA and MD were significantly correlated with these neurological changes (Figure 4B). In addition, FA displayed a significant correlation with motor related assessments, indicating the utility of FA in monitoring motor function changes. Behaviorally, the monkeys displayed an inability to use their right upper limb for tasks like retrieving food, relying instead on their unaffected left limb (Figure 2B). This compensatory behavior, indicative of altered motor function and spatial perception, is a well-documented response to stroke-induced impairments (Won et al., 2020) (Diomerick et al., 2006). Recent research indicates that compensatory use of the unaffected limb post-stroke activates the contralateral motor cortex, facilitating neural reorganization and functional recovery, as observed in enhanced motor cortex activity and functional connectivity (Jones et al., 2013) (Savidan et al.,2017). Cognitively, there was a noticeable decline in working memory as evidenced in the delayed-response task at 1-month post-MCAO (Figure 4B). This pattern, aligning with the changes in DTI metrics, points to the impact of brain damage on cognitive functions (Reijmer et al., 2013). However, the observed cognitive recovery by three months post-stroke highlights the brain’s resilience and capacity for recovery following such injuries.

Our MS/MS functional enrichment analysis identified two distinct groups, primarily involved in regulating cell morphogenesis and migration, apoptosis, and vascular functions (Figure 3). Of these, 15 proteins displayed a strong correlation with DTI metrics. Notably, MMP9, THBS1, MB and HSP90AA1 in Cluster 1 were negatively correlated with FA values, while SERPINB1, ACTBL2, ACTG1, Coro1A, RUS1, MGLL, WDR1, PFN1, TAGLN2, YWHAZ and ACTN1 in Cluster 2 showed a positive correlation with MD values. These proteins also correlated positively with neurological function scores and a negative correlation with DRT correct, suggesting an alignment between changes in behavioral phenotype, serum protein level, and brain pathological and functional alterations observed in MRI. MMP9, a key player in blood-brain barrier degradation, was consistently elevated in monkey serum up to 3 months post-MCAO. This sustained elevation aligns with clinical findings associating high MMP9 levels in ischemic stroke patients with increased risks of severe disability or death, underscoring its prognostic significance (Cui et al., 2012) (Turner & Sharp, 2016). Our findings also indicated a rise in THBS1 post-MCAO, a protein critical in hemostasis (Gao et al., 2015) and mediating cellular stress responses (Schips et al., 2019), mirroring human ischemic stroke findings where THBS1 elevation correlates with stroke severity (Qin et al., 2019). Elevated serum MB levels post-MCAO were observed, consistent with increase noted in human stroke patients (Di Angelantonio et al., 2005) and acute stroke monkey models (Guo et al., 2019). This sustained elevation aligns with clinical findings associating high MMP9 levels in ischemic stroke patients with increased risks of severe disability or death, underscoring its prognostic significance (Sato et al., 2005); Aboouf et al., 2023). PFN1, an actin-binding protein involved in various physiological processes, showed elevated levels post-MCAO. Present in almost all tissues and cells (Alkam et al., 2017), PFN1 is crucial for multiple physiological processes, including cell migration (Lin et al., 2018), vascular permeability (Li et al., 2013), angiogenesis (Fan & Fox, 2012) and oxidative stress (Li et al., 2018). Intriguingly, inhibiting PFN1 offered neuroprotection against ischemia/reperfusion injuries, partly by promoting M2 microglial polarization, indicating its potential as a therapeutic target in stroke management (Lu et al., 2020). YWHAZ, part of the 14-3-3 protein family, affects cellular proliferation, migration, and differentiation by interacting with phosphoserine-and serine-containing proteins (Aghazadeh & Papadopoulos, 2016). Clinically, elevated levels of YWHAZ post-stroke have been associated with cognitive impairment (Qi et al., 2023), while experimental models demonstrate that reduced YWHAZ expression leads to neurodevelopmental and cognitive deficits (Wan et al., 2023). Our study establishes the significance of five proteins in in ischemic stroke, highlighting their potential as biomarkers and therapeutic targets.

Our research identified 15 proteins correlated with DTI metrics, of which 10 proteins have not been previously reported in direct association with ischemic stroke. These proteins include HSP90AA1, SERPINB1, ACTBL2, ACTG1, Coro1A, RSU1, MGLL, WDR1, TAGLN2, ACTN1. While KEGG enrichment analyses showed their involvement in process like cell morphogenesis and migration, apoptosis and programmed cell death, and coagulation and blood vessel, their exact roles in the onset and recovery of ischemic stroke require further exploration. Although it is challenging for serum proteins to to mirror molecular changes in brain tissue, especially in non-human primate models due to difficulties in acquiring brain tissue for continuously detection, in-vivo monitoring combining serum protein, behavioral analysis and MRI imaging remains a valuable tool for long-term tracking of disease progression in animal models.

In summary, our study in a non-human primate model of ischemic stroke provides key insights into stroke’s complex dynamics, linking neuroimaging data with behavioral and proteomic changes. We highlighted the significance of certain serum proteins, including MMP9, THBS1, MB, PFN1, and YWHAZ, as potential biomarkers and therapeutic targets. Our integrative approach underscores the importance of combining neuroimaging, behavioral studies, and proteomics for a comprehensive understanding of stroke, offering promising avenues for enhancing stroke research and treatment strategies.

## Materials and Methods

### Animal preparation and surgery

Three male cynomolgus macaques (crab-eating macaque, 3050703, 13050217 and 13051415), aged 8.67 ± 0.57 years and weighting 8.38 ± 1.99 kg, were included in this study. Housed individually under a 12-h light/dark cycle, the monkeys were maintained in an environment controlled at 24±2°C with 40±20% humidity. They had free access to water and were fed twice daily. The middle cerebral artery occlusion (MCAO) model adapted from established protocols (Chen et al., 2015) was accomplished in the three monkeys using bipolar electrocoagulation. Comprehensive evaluations, including MRI data collection, behavior assessments, and serum proteomic analysis, were systematically conducted. These evaluations were performed at three timepoints: before the induction of MCAO to establish a baseline, and then at one month and three months following the MCAO procedure. All procedures adhered to the “Care and Use of Laboratory Animals” guidelines and were approved by the Institutional Animal Care and Use Committee of Guangdong Laboratory Animals Monitoring Institute (IACUC no. 2018014, AAALAC accredited).

For MCAO surgery, anesthesia was initiated with ketamine (10 mg/kg), xylazine (2mg/kg), and atropine (2mg/kg) intramuscularly, and maintained with 1–1.5% isoflurane vaporized in 100% oxygen. Vital signs, including heart rate, blood pressure, respiration, and oxygen saturation, were continuously monitored. The surgery involved aseptic removal of the left frontotemporal bone, exposure of the Sylvian fissure, and occlusion of the left MCA distal to the M1 branch. Post-surgery, the site was washed with saline, treated with bone wax, and sutured. Then, monkeys were housed in their cages with regulated body temperature for recovery. The first 3 days post-MCAO required thrice-daily hand-feeding due to unilateral impairments. Antibiotic (penicillin, 0.4 million units, intramuscularly) and mannitol (20%, intravenously) were administered twice daily for infection prevention and intracranial pressure reduction. A veterinarian monitored daily until independent self-care was re-established.

### MRI images acquisition and data processing

Using a 3.0 T MRI system (MAGNETOM Trio, Siemens Healthineers, Erlangen, Germany) equipped with an 8-channels head coil, we acquired MRI data from each anesthetized animal (maintained with 2.5% isoflurane). Animals were placed in an MR-compatible stereotactic apparatus for imaging. Structural images were obtained using a T2-weighted turbo spin echo sequence (TR = 3000 ms, TE = 396 ms, voxel size: 0.6 mm × 0.6 mm × 0.6 mm, and BW = 651 Hz). Accordingly, diffusion tensor imaging (DTI) was performed to provide microstructural images, using a multiband echo-planar sequence (TR = 5700 ms, TE = 99 ms, voxel size: 1.3 mm × 1.3 mm × 1.3 mm, and BW = 1502 Hz).

The infarct volume resulting from MCAO was manually outlined on T2-weighted images (T2WI) at 1-month and 3-month post-MCAO using ITK-SNAP software (version 3.8, http://www.itksnap.org). This software automatically calculated the infarct volume by summing the volume of identified lesions lesions. Regions of interest (ROIs) were defined on T2WI following the Paxinos Macaque Atlas (Calabrese et al., 2015) available at https://scalablebrainatlas.incf.org/macaque/CBCetal15. DTI data were processed with FSL (version 6.0, FMRIB Software Library, http://www.fmrib.ox.ac.uk/fsl) and DSI Studio (version 12, Yeh FC. https://dsi-studio.labsolver.org). DTI metrics, including fractional anisotropy (FA), mean diffusivity (MD), axial diffusivity (AD) and radical diffusivity (RD), were measured to evaluate microstructural properties and diffusion characteristics in the brain (Fan et al., 2016).

### Behavioral Assessment

This study conducted a comprehensive behavioral assessment in nonhuman primates, including neurologic deficit evaluation, motor function analysis (hill and valley staircase task, HVST), working memory assessment (delayed-response task, DRT), and cognitive function evaluation (object-retrieval detour task, ORDT). These assessments were conducted before MCAO induction, and at 1-month and 3-month post-MCAO.

Neurologic deficits were quantified using a standardized neurological deficit score (Kito et al., 2001), including four categories totaling 100 points: consciousness (28 points), sensory system (22 points), motor system (32 points), and skeletal muscle coordination (18 points). Two blinded, experienced observes performed these assessments, with a higher score indicating more server neurological damage.

We assessed motor function and spatial perception using HVST (Marshall & Ridley, 2003). Monkeys retrieved food rewards from staircases outside their cages, reaching through vertical slots in a front-panel plexiglass cage. Scores were assigned based on the distance from the slot (1 for nearest, 5 for furthest), with a maximum possible score of 15 per side. In the hill task, there are two staircases positioned such that they converge towards the center at the top. This arrangement resembles a hill. Conversely, in the valley task, the staircases are arranged to diverge from a central point, like a valley. Monkeys were given three minutes to retrieve all the food items (apple pieces) from the staircases. The order of testing was randomized to prevent bias. The maximum possible score of each staircase was 45.

Working memory was evaluated using DRT with the Wisconsin General Test Apparatus (Tsujimoto & Sawaguchi, 2002). Monkeys were trained on a two-well delayed response task before surgery. During the actual test, a piece of food was randomly and visibly placed in one of the two wells, and then both wells were covered with lids. An opaque clapboard as then placed between the monkey and wells, obstructing the view. After predetermined delay, the clapboard was removed, and the monkey was allowed to select one of the two lids to lift in search of the food reward. The final set of delays include 0, 4, 8, 12, and 16 seconds. The average correct response rate over five rounds indicated working memory proficiency.

Cognitive function was assessed using ORDT (Sutcliffe et al., 2014). Monkeys attempted to retrieve food from a transparent plexiglass box with varying open-side orientations. The test comprised 17 trials, with both easy (9 trials) and difficult (eight trails) food retrievals. Performance was quantified by success rate (first attempt success) and number of barrier hits (attempts to reach the reward through the transparent wall). Each monkey underwent five rounds with average success rate and barrier hits assessed.

### Proteomics

Blood samples were obtained from three animals before MCAO surgery and at 1-month and 3-month post-MCAO. Serum was extracted from these samples and stored at −80℃. Proteins in each serum sample were denatured, reduced, alkylated, and digested with sequencing-grade modified trypsin (protein-to-enzyme ratio of 50:1) at 37◦C overnight. Then, peptides were recovered through centrifugation (12000 g for 10 min) and desalted using Strata X SPE column.

Peptides, dissolved in solvent A, were loaded onto a homemade reversed-phase analytical column (25-cm length, 100 μm i.d.). A gradient consisting of solvent A (0.1% formic acid, 2% acetonitrile in water) and solvent B (0.1% formic acid, 90% acetonitrile in water) was employed for peptide separation on an ASY-nLC 1200 UPLC system (ThermoFisher Scientific). The separation gradient was set as follows: 0-68 min, 4%-20% B; 68-82 min, 20%-32% B; 82-86 min, 32%-80% B; 86-90 min, 80% B, maintaining a constant flow rate of 500 nl/min. The peptides were then analyzed using an Orbitrap Exploris 480 with a nano-electrospray ion source (electrospray voltage: 2300 V). FAIMS compensate voltage (CV) was set at -70 V, -45 V. The Oorbitrap detector were configured for optimal analysis with a full mass spectrometry (MS) scan resolution of 60,000 (scan range: 400-1200 m/z) and MS/MS scan fixed at a first mass of 110 m/z, resolution 30,000. TurboTMT was disabled. A maximum of 15 abundant precursors were selected for MS/MS analyses with a 30s dynamic exclusion. HCD fragmentation was performed at a normalized collision energy (NCE) of 27%, with an automatic gain control (AGC) target of 75%, an intensity threshold of 10,000 ions/s, and a maximum injection time of 100 ms. The resulting MS/MS data were processed using the Proteome Discoverer search engine (v.2.4), searching against the Macaca_fascicularis_9541_PR_20220314.fasta database (50208 entries). Search parameters included Trypsin (Full) as the cleavage enzyme, allowing for up to 2 missed cleavages, a minimum peptide length of 6, and a maximum of a modifications per peptide. Mass error setting were 10 ppm for precursor iron and 0.02 Da for fragment ions. Fixed modification was Carbamidomethyl on Cys, while variable modifications included Oxidation on Met, Acetylation on protein N-terminal, met-loss on Met, and met-loss+acetyl on Met. To false discovery rate (FDR) was adjusted to less than 1% for protein, peptide, and PSM levels to ensure high data confidence.

For gene ontology and pathway analysis, the org.Hs.eg.db R package from Bioconductor was used to correlate UniProtKB accession and gene name with ENTREZ Gene IDs. Functional classification and enrichment analyses of gene ontology (GO) terms among differentially expressed proteins were performed using ENTREZ Gene IDs via the clusterProfiler R package. Kyoto Encyclopedia of Genes and Genomes (KEGG) pathway enrichment analysis was also conducted using clusterProfiler R package, supported by a bioconductor package for pathway-based data integration and visualization.

### Statistical Analysis

Statistical analyses were conducted using GraphPad Prism (version 9, GraphPad Inc., San Diego, CA, USA). Data were represented as the Mean ± Standard Error of the Mean (SEM). A two-sample t-test was employed to compare ipsilateral and contralateral changes at baseline (before MCAO), and at 1-month and 3-month post-MCAO timepoints. Difference across the three timepoints (aseline, 1-month, and 3-month post-MCAO) were analyzed using two-way ANOVA. was carried out with three time points, followed by *post hoc* analysis using Tukey’s test was applied for post hoc analysis to identify specific group differences. The relationships between MRI data, behavioral outcomes, and serum proteomics were explored using Spearman correlation coefficients. To integrate multiple biomarkers, including DTI metrics, behavioral scores, and serum proteomics, simple linear regression analysis was employed. This approach was intended to provide a more comprehensive understanding of the interplay between different types of data.

### Author Information

#### Corresponding Author

**Hsin-Yi Lai** - Department of Neurology of the Second Affiliated Hospital, Interdisciplinary Institute of Neuroscience and Technology, Zhejiang University School of Medicine, Hangzhou, 310029, China; College of Biomedical Engineering and Instrument Science, Zhejiang University, Hangzhou, 310027, China; Liangzhu Laboratory, MOE Frontier Science Center for Brain Science and Brain-Machine Integration, State Key Laboratory of Brain-machine Intelligence, School of Brain Science and Brain Medicine, Zhejiang University, Hangzhou, 310058, China; Affiliated Mental Health Center & Hangzhou Seventh People’s Hospital, Zhejiang University School of Medicine, Zhejiang University, Hangzhou, 310013, China; **Yu Zhang** - Guangdong Provincial Key Laboratory of Laboratory Animals, Guangdong Laboratory Animals Monitoring Institute, Guangzhou, 510663, China;

#### Authors

**Ge Li** - Guangdong Provincial Key Laboratory of Laboratory Animals, Guangdong Laboratory Animals Monitoring Institute, Guangzhou, 510663, China

**Lan Lan** - Department of Neurology of the Second Affiliated Hospital, Interdisciplinary Institute of Neuroscience and Technology, Zhejiang University School of Medicine, Hangzhou, 310029, China; Department of Psychology and Behavior Science, Zhejiang University, Hangzhou, 310028, China

**Tingting He** - Department of Neurology of the Second Affiliated Hospital, Interdisciplinary Institute of Neuroscience and Technology, Zhejiang University School of Medicine, Hangzhou, 310029, China; College of Biomedical Engineering and Instrument Science, Zhejiang University, Hangzhou, 310027, China

**Zheng Tang** - Department of Neurology of the Second Affiliated Hospital, Interdisciplinary Institute of Neuroscience and Technology, Zhejiang University School of Medicine, Hangzhou, 310029, China

**Shuhua Liu** - Guangdong Provincial Key Laboratory of Laboratory Animals, Guangdong Laboratory Animals Monitoring Institute, Guangzhou, 510663, China

**Yunfeng Li** - Guangdong Provincial Key Laboratory of Laboratory Animals, Guangdong Laboratory Animals Monitoring Institute, Guangzhou, 510663, China

**Zhongqiang Huang** - Guangdong Provincial Key Laboratory of Laboratory Animals, Guangdong Laboratory Animals Monitoring Institute, Guangzhou, 510663, China

**Yalun Guan** - Guangdong Provincial Key Laboratory of Laboratory Animals, Guangdong Laboratory Animals Monitoring Institute, Guangzhou, 510663, China

**Xuejiao Li** - Guangdong Provincial Key Laboratory of Laboratory Animals, Guangdong Laboratory Animals Monitoring Institute, Guangzhou, 510663, China

#### Author Contributions

Conceptualization: H.Y.L., Y.Z.

Methodology: G.L., L.L., T.H., Z.T.

Investigation: G.L., S.L., Y.L., Z.H., Y.G., X.L.

Visualization: L.L.

Supervision: H.Y.L., Y.Z.

Writing original draft: L.L., G.L.

Writing review & editing: H.Y.L., Y.Z.

### Notes

Ethics Approval. Animals were used after approval from the Institutional Animal Care and Use Committee of Guangdong Laboratory Animals Monitoring Institute (IACUC no. 2018014, AAALAC accredited).

The authors declare no competing financial interest.

Data and materials availability: All data are available in the main text or the supplementary materials.

## Acknowledgements

This work was supported by National Key R&D Program of China (2021YFF0702200), Guangdong provincial science and technology plan project (2009A081000002, 2008A08003), National Natural Science Foundation of China (82101323), Key R&D Program of Zhejiang Province (2021C03001), Guangzhou Science and technology planning project (202206010084)

## References

1. Aboouf, M. A., Gorr, T. A., Hamdy, N. M., Gassmann, M., & Thiersch, M. (2023). Myoglobin in Brown Adipose Tissue: A Multifaceted Player in Thermogenesis. Cells, 12(18), 2240.

2. Aghazadeh, Y., & Papadopoulos, V. (2016). The role of the 14-3-3 protein family in health, disease, and drug development. Drug Discov. Today, 21(2), 278–287.

3. Alkam, D., Feldman, E. Z., Singh, A., & Kiaei, M. (2017). Profilin1 biology and its mutation, actin(g) in disease. Cell. Mol. Life Sci., 74(6), 967–981.

4. Bai, Y. L., Hu, Y. S., Wu, Y., Zhu, Y. L., Zhang, B., Jiang, C. Y., … & Fan, W.K. (2014). Long-term three-stage rehabilitation intervention alleviates spasticity of the elbows, fingers, and plantar flexors and improves activities of daily living in ischemic stroke patients: a randomized, controlled trial. Neuroreport, 25(13), 998–1005.

5. Chen, L., Peters, J. E., Prins, B., Persyn, E., Traylor, M., Surendran, P., … & Howson, J.M. (2022). Systematic Mendelian randomization using the human plasma proteome to discover potential therapeutic targets for stroke. Nat. Commun., 13(1), 6143.

6. Chen, X., Dang, G., Dang, C., Liu, G., Xing, S., Chen, Y., … & Zeng, J. (2015). An ischemic stroke model of nonhuman primates for remote lesion studies: a behavioral and neuroimaging investigation. Restor. Neurol. Neurosci., 33(2), 131–142.

7. Cui, J., Chen, S., Zhang, C., Meng, F., Wu, W., Hu, R., … & Gu, Z. (2012). Inhibition of MMP-9 by a selective gelatinase inhibitor protects neurovasculature from embolic focal cerebral ischemia. Mol. Neurodegener., 7(1), 1–15.

8. Di Angelantonio, E., Fiorelli, M., Toni, D., Sacchetti, M. L., Lorenzano, S., Falcou, A., Ciarla, M. V., Suppa, M., Bonanni, L., Bertazzoni, G., Aguglia, F., & Argentino, C. (2005). Prognostic significance of admission levels of troponin I in patients with acute ischaemic stroke. J. Neurol. Neurosurg. Psychiatry, 76(1), 76–81.

9. Dromerick, A. W., Lang, C. E., Birkenmeier, R., Hahn, M. G., Sahrmann, S. A., & Edwards, D. F. (2006). Relationships between upper-limb functional limitation and self-reported disability 3 months after stroke. J. Rehabil. Res. Dev., 43(3), 401–408.

10. Fan, S., Van den Heuvel, O. A., Cath, D. C., Van der Werf, Y. D., De Wit, S. J., De Vries, F.E., … & Pouwels, P.J. (2016). Mild white matter changes in un-medicated obsessive-compulsive disorder patients and their unaffected siblings. Front. Neurosci., 9, 495.

11. Fan, Y., & Fox, P. L. (2012). Alternative angiogenic pathway driven by stimulus-dependent phosphorylation of profilin-1. Nat. Cell Biol., 14(10), 1046–1056.

12. Fisher, M., Feuerstein, G., Howells, D. W., Hurn, P. D., Kent, T. A., Savitz, S. I., & Lo, E. H. (2009). Update of the stroke therapy academic industry roundtable preclinical recommendations. Stroke, 40(6), 2244-2250.

13. Foltys, H., Krings, T., Meister, I. G., Sparing, R., Boroojerdi, B., Thron, A., & Töpper, R. (2003). Motor representation in patients rapidly recovering after stroke: a functional magnetic resonance imaging and transcranial magnetic stimulation study. Clin. Neurophysiol., 114(12), 2404–2415.

14. Gao, J. B., Tang, W. D., Wang, H. X., & Xu, Y. (2015). Predictive value of thrombospondin-1 for outcomes in patients with acute ischemic stroke. Clin. Chim. Acta, 450, 176–180.

15. GBD 2016 Lifetime Risk of Stroke Collaborators. (2018). Global, regional, and country-specific lifetime risks of stroke, 1990 and 2016. N. Engl. J. Med., 379(25), 2429-2437.

16. Guo, L., Zhou, D. A., Wu, D. I., Ding, J., He, X., Shi, J., … & Meng, R. (2019). ShortCterm remote ischemic conditioning may protect monkeys after ischemic stroke. Ann. Clin. Transl. Neurol., 6(2), 310–323.

17. Hagmann, P., Jonasson, L., Maeder, P., Thiran, J. P., Wedeen, V. J., & Meuli, R. (2006). Understanding diffusion MR imaging techniques: from scalar diffusion-weighted imaging to diffusion tensor imaging and beyond. Radiographics, 26(suppl_1), S205-S223.

18. Hochrainer, K., & Yang, W. (2022). Stroke proteomics: from discovery to diagnostic and therapeutic applications. Circ. Res., 130(8), 1145–1166.

19. Jones, T. A. (2017). Motor compensation and its effects on neural reorganization after stroke. Nat. Rev. Neurosci., 18(5), 267–280.

20. Jones, T. A., Allred, R. P., Jefferson, S. C., Kerr, A. L., Woodie, D. A., Cheng, S. Y., & Adkins, D. L. (2013). Motor system plasticity in stroke models: intrinsically use-dependent, unreliably useful. Stroke, 44(6_suppl_1), S104-S106.

21. Jung, W. B., Han, Y. H., Chung, J. J., Chae, S. Y., Lee, S. H., Im, G. H., … & Lee, J.H. (2017). Spatiotemporal microstructural white matter changes in diffusion tensor imaging after transient focal ischemic stroke in rats. NMR Biomed., 30(6), e3704.

22. Kito, G., Nishimura, A., Susumu, T., Nagata, R., Kuge, Y., Yokota, C., & Minematsu, K. (2001). Experimental thromboembolic stroke in cynomolgus monkey. J. Neurosci. Methods, 105(1), 45–53.

23. Kronlage, M., Schwehr, V., Schwarz, D., Godel, T., Uhlmann, L., Heiland, S., … & Bäumer, P. (2018). Peripheral nerve diffusion tensor imaging (DTI): normal values and demographic determinants in a cohort of 60 healthy individuals. Eur. Radiol., 28, 1801-1808.

24. Kwak, S. Y., Yeo, S. S., Choi, B. Y., Chang, C. H., & Jang, S. H. (2010). Corticospinal tract change in the unaffected hemisphere at the early stage of intracerebral hemorrhage: a diffusion tensor tractography study. Eur. Neurol., 63(3), 149–153.

25. Langhorne, P., Coupar, F., & Pollock, A. (2009). Motor recovery after stroke: a systematic review. Lancet Neurol., 8(8), 741–754.

26. Li, C. X., Meng, Y., Yan, Y., Kempf, D., Howell, L., Tong, F., & Zhang, X. (2022). Investigation of white matter and grey matter alteration in the monkey brain following ischemic stroke by using diffusion tensor imaging. Invest. Magn. Reson. Imaging, 26(4), 275.

27. Li, X., Liu, J., Chen, B., & Fan, L. (2018). A positive feedback loop of profilin-1 and RhoA/ROCK1 promotes endothelial dysfunction and oxidative stress. Oxid. Med. Cell. Longev., 2018.

28. Li, Z., Zhong, Q., Yang, T., Xie, X., & Chen, M. (2013). The role of profilin-1 in endothelial cell injury induced by advanced glycation end products (AGEs). Cardiovasc Diabetol, 12(1), 1–11.

29. Lin, W., Izu, Y., Smriti, A., Kawasaki, M., Pawaputanon, C., Böttcher, R. T., … & Ezura, Y. (2018). Profilin1 is expressed in osteocytes and regulates cell shape and migration. J. Cell. Physiol., 233(1), 259–268.

30. Lindenberg, R., Zhu, L. L., Rüber, T., & Schlaug, G. (2012). Predicting functional motor potential in chronic stroke patients using diffusion tensor imaging. Hum. Brain Mapp., 33(5), 1040–1051.

31. Lu, E., Wang, Q., Li, S., Chen, C., Wu, W., Xu, Y. X. Z., … & Zhou, K. (2020). Profilin 1 knockdown prevents ischemic brain damage by promoting M2 microglial polarization associated with the RhoA/ROCK pathway. J. Neurosci. Res., 98(6), 1198–1212.

32. Ma, C., Liu, A., Li, Z., Zhou, X., & Zhou, S. (2014). Longitudinal study of diffusion tensor imaging properties of affected cortical spinal tracts in acute and chronic hemorrhagic stroke. J. Clin. Neurosci., 21(8), 1388–1392.

33. Marshall, J. W., & Ridley, R. M. (2003). Assessment of cognitive and motor deficits in a marmoset model of stroke. ILAR J., 44(2), 153–160.

34. Marshall, J. W., Duffin, K. J., Green, A. R., & Ridley, R. M. (2001). NXY-059, a free radical– trapping agent, substantially lessens the functional disability resulting from cerebral ischemia in a primate species. Stroke, 32(1), 190–198.

35. Narayan, S. K., Grace Cherian, S., Babu Phaniti, P., Babu Chidambaram, S., Rachel Vasanthi, A. H., & Arumugam, M. (2021). Preclinical animal studies in ischemic stroke: Challenges and some solutions. Anim. Models Exp. Med., 4(2), 104–115.

36. Okamoto, Y., Ishii, D., Yamamoto, S., Ishibashi, K., Wakatabi, M., Kohno, Y., & Numata, K. (2021). Relationship between motor function, DTI, and neurophysiological parameters in patients with stroke in the recovery rehabilitation unit. J. Stroke Cerebrovasc. Dis., 30(8), 105889.

37. Pasi, M., van Uden, I. W., Tuladhar, A. M., de Leeuw, F. E., & Pantoni, L. (2016). White matter microstructural damage on diffusion tensor imaging in cerebral small vessel disease: clinical consequences. Stroke, 47(6), 1679–1684.

38. Qi, B., Kong, L., Lai, X., Wang, L., Liu, F., Ji, W., & Wei, D. (2023). Plasma exosome proteomics reveals the pathogenesis mechanism of post-stroke cognitive impairment. Aging (Albany NY), 15(10), 4334.

39. Qin, C., Zhao, X. L., Ma, X. T., Zhou, L. Q., Wu, L. J., Shang, K., … & Tian, D.S. (2019). Proteomic profiling of plasma biomarkers in acute ischemic stroke due to large vessel occlusion. J. Transl. Med., 17, 1–9.

40. Reijmer, Y. D., Freeze, W. M., Leemans, A., & Biessels, G. J. (2013). The effect of lacunar infarcts on white matter tract integrity. Stroke, 44(7), 2019–2021.

41. Sato, Y., Iwamoto, J., Kanoko, T., & Satoh, K. (2005). Negative myoglobin staining in hemiplegic muscle of acute stroke patients predicts functional recovery. Am. J. Phys. Med. Rehabil., 84(9), 692–698.

42. Schips, T. G., Vanhoutte, D., Vo, A., Correll, R. N., Brody, M. J., Khalil, H., … & Molkentin, J.D. (2019). Thrombospondin-3 augments injury-induced cardiomyopathy by intracellular integrin inhibition and sarcolemmal instability. Nat. Commun., 10(1), 76.

43. Shaheen, H. A., Sayed, S. S., Magdy, M. M., Saad, M. A., Magdy, A. M., & Daker, L. I. (2022). Prediction of motor recovery after ischemic stroke: Clinical and diffusion tensor imaging study. J Clin Neurosci, 96, 68–73.

44. Shen, T., Yue, Y., Ba, F., He, T., Tang, X., Hu, X., … & Lai, H.Y. (2022). Diffusion along perivascular spaces as marker for impairment of glymphatic system in Parkinson’s disease. NPJ Parkinson’s Dis, 8(1), 174.

45. Sironi, L., Tremoli, E., Miller, I., Guerrini, U., Calvio, A. M., Eberini, I., … & Gianazza, E. (2001). Acute-phase proteins before cerebral ischemia in stroke-prone rats: identification by proteomics. Stroke, 32(3), 753–760.

46. Sommer, C. J. (2017). Ischemic stroke: experimental models and reality. Acta Neuropathol, 133(2), 245–261.

47. Stevens, S. L., Liu, T., Bahjat, F. R., Petyuk, V. A., Schepmoes, A. A., Sontag, R. L., … & Stenzel-Poore, M. P. (2019). Preconditioning in the rhesus macaque induces a proteomic signature following cerebral ischemia that is associated with neuroprotection. Transl Stroke Res, 10, 440–448.

48. Sutcliffe, J. S., Beaumont, V., Watson, J. M., Chew, C. S., Beconi, M., Hutcheson, D. M., … & Munoz-Sanjuan, I. (2014). Efficacy of selective PDE4D negative allosteric modulators in the object retrieval task in female cynomolgus monkeys (Macaca fascicularis). PLoS One, 9(7), e102449.

49. Taha, A., Bobi, J., Dammers, R., Dijkhuizen, R. M., Dreyer, A. Y., van Es, A. C., … & van Beusekom, H. M. (2022). Comparison of large animal models for acute ischemic stroke: which model to use?. Stroke, 53(4), 1411–1422.

50. Tsujimoto, S., & Sawaguchi, T. (2002). Working memory of action: a comparative study of ability to selecting response based on previous action in New World monkeys (Saimiri sciureus and Callithrix jacchus). Behav Processes, 58(3), 149–155.

51. Turner, R. J., & Sharp, F. R. (2016). Implications of MMP9 for blood brain barrier disruption and hemorrhagic transformation following ischemic stroke. Front Cell Neurosci, 10, 56.

52. Wan, R. P., Liu, Z. G., Huang, X. F., Kwan, P., Li, Y. P., Qu, X. C., … & Qiao, J.D. (2023). YWHAZ variation causes intellectual disability and global developmental delay with brain malformation. Hum Mol Genet, 32(3), 462–472.

53. Won, J., Yi, K. S., Choi, C. H., Jeon, C. Y., Seo, J., Kim, K., … & Lee, Y. (2020). Assessment of hand motor function in a non-human primate model of ischemic stroke. Exp Neurol, 29(4), 300.

54. Yi, K. S., Choi, C. H., Lee, S. R., Lee, H. J., Lee, Y., Jeong, K. J., … & Cha, S.H. (2017). Sustained diffusion reversal with in-bore reperfusion in monkey stroke models: confirmed by prospective magnetic resonance imaging. J Cereb Blood Flow Metab, 37(6), 2002–2012.

55. Zweier, J. L., & Talukder, M. H. (2006). The role of oxidants and free radicals in reperfusion injury. Cardiovasc Res, 70(2), 181–190.

