## Supplementary material for "Comprehensive Assessment of Ischemic Stroke in Nonhuman Primates: Neuroimaging, Behavioral, and Serum Proteomic Analysis": Table S1

**Supplemental Materials**

We present our findings on axial diffusivity (AD) and radial diffusivity (RD) values, further characterizing microstructural changes in the brain post-stroke, as shown in Figure S1. Notably, we observed significant disparities in AD values within the caudate nucleus (Cd) on the ipsilateral side when compared to the contralateral regions. Additionally, by the 3-month post-MCAO mark, increased AD values were evident across several ipsilateral regions of interest (ROIs), including the Cd, putamen (Pu), orbitofrontal cortex (OC), and insular cortex (IC). RD values, on the other hand, displayed an upward trend over the course of the 3-month post-MCAO period. A significant elevation in RD values was observed in the ipsilateral ROIs compared to their contralateral counterparts, except for the olfactory tubercle (OT). Intriguingly, the OT exhibited a decrease in RD at 1-month post-MCAO, followed by an increase from 1-month to 3-month post-MCAO.

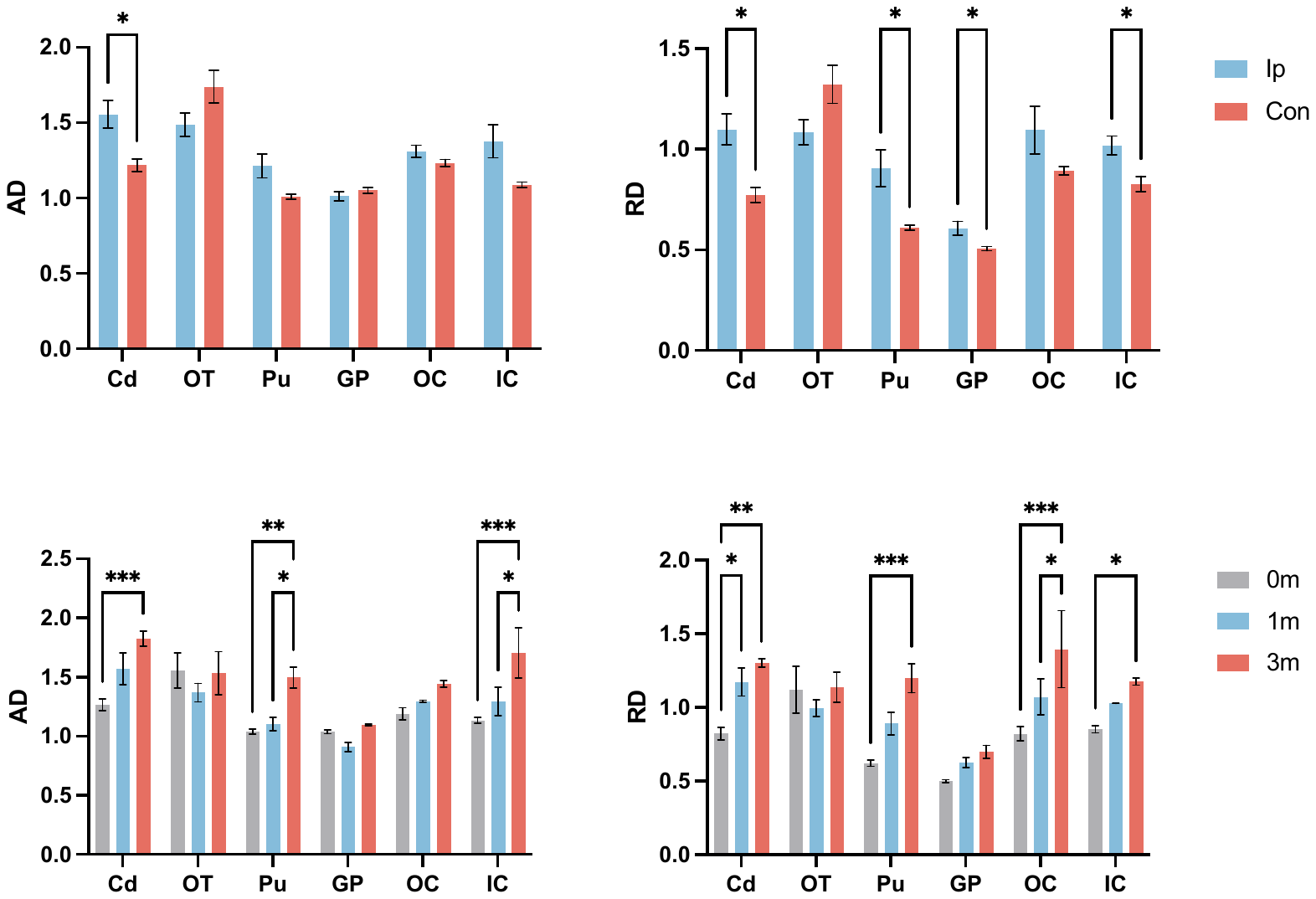

**Fig. S1** Comparison of AD and RD values between ipsilateral and contralateral sides across six ROIs. Comparison of AD and RD values of the ipsilateral side across six ROIs at baseline, 1-month, and 3-month. Significance levels are denoted as follows: ***, P<0.001. **, P<0.01. *, P<0.05**.** Cd, Caudate Nucleus; OT, Olfactory Tubercle; Pu, Putamen; GP, Globus Pallidus; OC, Orbitofrontal cortex; IC, Insular Cortex; Con, contralateral; IP, ipsilateral. Data are represented as the Mean ± SEM.

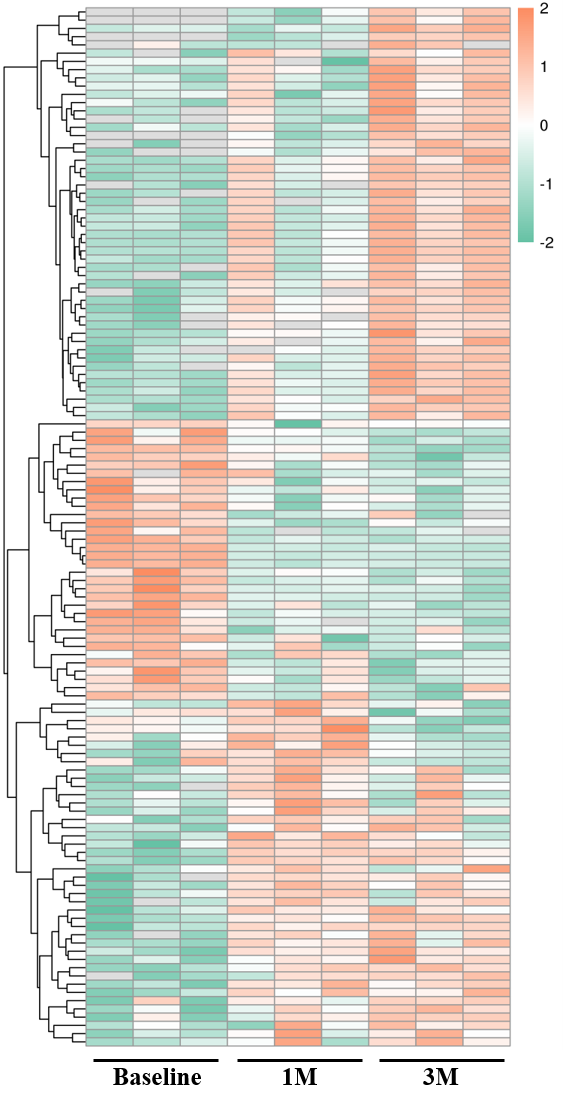

**Fig. S2** Hierarchical cluster analysis (HCA) of 5214 proteins.

**Table S1** Candidate proteins

| Protein accession | Protein description | Gene name | 1M/Baseline Ratio | 1M/Baseline P value | 3M/Baseline Ratio | 3M/Baseline P value | 3M/1M Ratio | 3M/1M P value | Mfuzz Cluster |
| --- | --- | --- | --- | --- | --- | --- | --- | --- | --- |
| A0A2K5TWD9 | Multimerin 1 OS=Macaca fascicularis OX=9541 GN=MMRN1 PE=4 SV=2 | MMRN1 | 36.167 | 0.010234666 | 32.167 | 0.022044206 | 0.889 | 0.551555466 | Cluster1 |
| A0A2K5UU71 | 92 kDa gelatinase OS=Macaca fascicularis OX=9541 GN=MMP9 PE=3 SV=2 | MMP9 | 41.975 | 0.000338906 | 32.252 | 0.001279534 | 0.768 | 0.397997045 | Cluster1 |
| A0A2K5V619 | Thrombospondin 1 OS=Macaca fascicularis OX=9541 PE=3 SV=2 | -- | 17.076 | 0.016979078 | 21.744 | 0.009988067 | 1.273 | 0.327981119 | Cluster1 |
| A0A2K5W341 | Vasodilator stimulated phosphoprotein OS=Macaca fascicularis OX=9541 GN=VASP PE=3 SV=2 | VASP | 20.368 | 0.114110091 | 15.231 | 0.037576296 | 0.748 | 0.861376303 | Cluster1 |
| P02150 | Myoglobin OS=Macaca fascicularis OX=9541 GN=MB PE=1 SV=2 | MB | 8.427 | 0.015511048 | 4.851 | 0.056365582 | 0.576 | 0.281434266 | Cluster1 |
| Q4R4P1 | Heat shock protein HSP 90-alpha OS=Macaca fascicularis OX=9541 GN=HSP90AA1 PE=2 SV=3 | HSP90AA1 | 7.667 | 0.282680482 | 8.213 | 0.015618026 | 1.071 | 0.526599577 | Cluster1 |
| A0A2K5TM01 | HIT domain-containing protein OS=Macaca fascicularis OX=9541 PE=3 SV=1 | -- | 3.398 | 0.032536894 | 11.943 | 0.00045223 | 3.514 | 0.022360358 | Cluster2 |
| A0A2K5TTB9 | Serpin family B member 1 OS=Macaca fascicularis OX=9541 GN=SERPINB1 PE=3 SV=2 | SERPINB1 | 3.873 | 0.005995589 | 7.522 | 0.001594302 | 1.942 | 0.014489905 | Cluster2 |
| A0A2K5TWA8 | Glyceraldehyde-3-phosphate dehydrogenase OS=Macaca fascicularis OX=9541 PE=3 SV=1 | -- | 2.42 | 0.045918511 | 7.634 | 0.001926745 | 3.155 | 0.035933314 | Cluster2 |
| A0A2K5UBM4 | Peptidyl-prolyl cis-trans isomerase OS=Macaca fascicularis OX=9541 PE=3 SV=2 | -- | 14.608 | 0.014373352 | 39.835 | 9.01042E-05 | 2.727 | 0.099261291 | Cluster2 |
| A0A2K5UGI0 | Actin beta like 2 OS=Macaca fascicularis OX=9541 GN=ACTBL2 PE=3 SV=2 | ACTBL2 | 7.82 | 0.105409042 | 21.122 | 0.000303788 | 2.701 | 0.133090991 | Cluster2 |
| A0A2K5UNU0 | X-prolyl aminopeptidase 1 OS=Macaca fascicularis OX=9541 GN=XPNPEP1 PE=3 SV=2 | XPNPEP1 | 6.33 | 0.139087162 | 10.993 | 0.007841893 | 1.737 | 0.349264522 | Cluster2 |
| A0A2K5UQ63 | Actin gamma 1 OS=Macaca fascicularis OX=9541 GN=ACTG1 PE=3 SV=2 | ACTG1 | 3.411 | 0.129918496 | 10.993 | 0.000237719 | 3.223 | 0.063010707 | Cluster2 |
| A0A2K5US25 | Coronin OS=Macaca fascicularis OX=9541 GN=CORO1A PE=3 SV=2 | CORO1A | 5.294 | 0.024253802 | 13.672 | 0.001831306 | 2.582 | 0.045026735 | Cluster2 |
| A0A2K5UTN5 | SH3 domain binding glutamate rich protein like 3 OS=Macaca fascicularis OX=9541 GN=SH3BGRL3 PE=3 SV=1 | SH3BGRL3 | 5.626 | 0.031809876 | 11.805 | 0.004807501 | 2.098 | 0.059748484 | Cluster2 |
| A0A2K5UVC3 | Ras suppressor protein 1 OS=Macaca fascicularis OX=9541 GN=RSU1 PE=4 SV=2 | RSU1 | 18.324 | 0.030387132 | 61.778 | 0.006382633 | 3.371 | 0.133726315 | Cluster2 |
| A0A2K5V664 | Monoglyceride lipase OS=Macaca fascicularis OX=9541 GN=MGLL PE=4 SV=2 | MGLL | 6.558 | 0.021269971 | 19.124 | 0.004615816 | 2.916 | 0.177765247 | Cluster2 |
| A0A2K5VU17 | Profilin OS=Macaca fascicularis OX=9541 GN=PFN1 PE=3 SV=1 | PFN1 | 20.368 | 0.037323891 | 70.284 | 0.00789449 | 3.451 | 0.079406737 | Cluster2 |
| A0A2K5W434 | Four and a half LIM domains 1 OS=Macaca fascicularis OX=9541 GN=FHL1 PE=4 SV=1 | FHL1 | 22.769 | 0.150124651 | 76.513 | 0.014231489 | 3.36 | 0.17108043 | Cluster2 |
| A0A2K5W8F0 | WD repeat domain 1 OS=Macaca fascicularis OX=9541 GN=WDR1 PE=4 SV=1 | WDR1 | 6.629 | 0.229060464 | 30.568 | 0.001938591 | 4.611 | 0.081347483 | Cluster2 |
| A0A2K5WBB8 | Protein S100 OS=Macaca fascicularis OX=9541 GN=EGM_01112 PE=3 SV=2 | EGM_01112 | 6.586 | 0.184088063 | 15.33 | 0.003510405 | 2.328 | 0.15949343 | Cluster2 |
| A0A2K5WCN1 | Moesin OS=Macaca fascicularis OX=9541 GN=MSN PE=4 SV=2 | MSN | 14.655 | 0.092986656 | 31.55 | 0.003329886 | 2.153 | 0.226054895 | Cluster2 |
| A0A2K5WDT3 | Parvin beta OS=Macaca fascicularis OX=9541 GN=PARVB PE=3 SV=2 | PARVB | 3.094 | 0.079183885 | 9.292 | 0.012515888 | 3.003 | 0.186838095 | Cluster2 |
| A0A2K5WK68 | Coronin OS=Macaca fascicularis OX=9541 GN=CORO1B PE=3 SV=2 | CORO1B | 4.481 | 0.110306046 | 14.597 | 0.000605544 | 3.258 | 0.079476747 | Cluster2 |
| A0A2K5WVG2 | F-actin-capping protein subunit alpha OS=Macaca fascicularis OX=9541 GN=CAPZA2 PE=3 SV=2 | CAPZA2 | 45.111 | 0.278169992 | 127.4 | 0.010560063 | 2.824 | 0.217689994 | Cluster2 |
| A0A2K5WZ75 | Transgelin OS=Macaca fascicularis OX=9541 GN=TAGLN2 PE=3 SV=1 | TAGLN2 | 2.253 | 0.17747505 | 5.214 | 0.003763158 | 2.314 | 0.10295246 | Cluster2 |
| A0A7N9CKG2 | L-lactate dehydrogenase OS=Macaca fascicularis OX=9541 PE=3 SV=1 | -- | 5.884 | 0.000927504 | 9.698 | 0.02698593 | 1.648 | 0.684097586 | Cluster2 |
| A0A7N9D1Q3 | Coactosin like F-actin binding protein 1 OS=Macaca fascicularis OX=9541 PE=4 SV=1 | -- | 28.143 | 0.120167756 | 95.529 | 0.000435713 | 3.394 | 0.15336622 | Cluster2 |
| A0A7N9IDK3 | Uncharacterized protein OS=Macaca fascicularis OX=9541 PE=4 SV=1 | -- | 6.045 | 0.027689938 | 16.508 | 0.003003452 | 2.731 | 0.098756233 | Cluster2 |
| G7PCF6 | Tyrosine 3-monooxygenase/tryptophan 5-monooxygenase activation protein zeta OS=Macaca fascicularis OX=9541 GN=YWHAZ PE=3 SV=1 | YWHAZ | 2.228 | 0.083094795 | 5.149 | 0.002409354 | 2.311 | 0.052112331 | Cluster2 |
| G7PFR8 | Tubulin beta chain OS=Macaca fascicularis OX=9541 GN=TUBB1 PE=3 SV=1 | TUBB1 | 7.194 | 0.195358594 | 22.434 | 0.005599594 | 3.118 | 0.142308496 | Cluster2 |
| G7PP69 | GST class-pi (Fragment) OS=Macaca fascicularis OX=9541 GN=EGM_05360 PE=3 SV=1 | EGM_05360 | 4.059 | 0.243081364 | 13.69 | 0.011503562 | 3.373 | 0.091881931 | Cluster2 |
| I7G4Q2 | Macaca fascicularis brain cDNA clone: QtrA-18223, similar to human tropomyosin 4 (TPM4), mRNA, RefSeq: NM_003290.1 OS=Macaca fascicularis OX=9541 PE=2 SV=1 | -- | 20.244 | 0.023571806 | 84.634 | 0.000226604 | 4.181 | 0.063576707 | Cluster2 |
| Q2PFV7 | Alpha-actinin-1 OS=Macaca fascicularis OX=9541 GN=ACTN1 PE=2 SV=1 | ACTN1 | 5.31 | 0.162000753 | 21.958 | 0.006839282 | 4.135 | 0.065267352 | Cluster2 |
| A0A2K5UDV1 | Nucleoside diphosphate kinase OS=Macaca fascicularis OX=9541 GN=NME1 PE=3 SV=2 | NME1 | 3.302 | 0.184020147 | 26.231 | 0.005089557 | 7.944 | 0.012710253 | Cluster4 |
| A0A2K5V4F1 | Triosephosphate isomerase OS=Macaca fascicularis OX=9541 PE=3 SV=1 | -- | 0.859 | 0.507961993 | 7.492 | 0.001484013 | 8.727 | 0.003512011 | Cluster4 |
| A0A2K5VBY5 | Filamin A OS=Macaca fascicularis OX=9541 GN=FLNA PE=3 SV=2 | FLNA | 1.522 | 0.573753313 | 5.217 | 0.00688027 | 3.427 | 0.043636187 | Cluster4 |
| A0A2K5TY00 | Fibrinogen gamma chain OS=Macaca fascicularis OX=9541 GN=FGG PE=4 SV=2 | FGG | 0.069 | 0.000134283 | 0.071 | 1.99287E-05 | 1.028 | 0.794468021 | Cluster5 |
| A0A2K5VSR8 | Fibrinogen alpha chain OS=Macaca fascicularis OX=9541 PE=4 SV=1 | -- | 0.068 | 1.40849E-05 | 0.062 | 3.40112E-06 | 0.913 | 0.313978681 | Cluster5 |
| G7P6F7 | Fibrinogen beta chain OS=Macaca fascicularis OX=9541 GN=FGB PE=4 SV=1 | FGB | 0.004 | 2.02953E-06 | 0.004 | 7.54583E-05 | 0.915 | 0.600768063 | Cluster5 |
| P68109 | Fibrinogen alpha chain (Fragment) OS=Macaca fascicularis OX=9541 GN=FGA PE=1 SV=1 | FGA | 0.128 | 0.000262492 | 0.015 | 0.000119923 | 0.115 | 0.002526614 | Cluster5 |
| A0A2K5VK62 | Carbamoyl-phosphate synthase 1 OS=Macaca fascicularis OX=9541 GN=CPS1 PE=3 SV=1 | CPS1 | 0.065 | 0.006205646 | 0.451 | 0.278768787 | 6.989 | 0.663070134 | Cluster6 |
